# The histone demethylase KDM5 controls developmental timing in *Drosophila* by promoting prothoracic gland endocycles

**DOI:** 10.1101/617985

**Authors:** Coralie Drelon, Helen M. Belalcazar, Julie Secombe

## Abstract

In *Drosophila*, the larval prothoracic gland integrates nutritional status with developmental signals to regulate growth and maturation through the secretion of the steroid hormone ecdysone. While the nutritional signals and cellular pathways that regulate prothoracic gland function are relatively well studied, the transcriptional regulators that orchestrate the activity of this tissue remain largely unknown. Here we show that <u>lysine d</u>e<u>m</u>ethylase 5 (KDM5) is essential for prothoracic gland function. Indeed, restoring *kdm5* expression only in the prothoracic gland in an otherwise *kdm5* mutant animal is sufficient to rescue both the larval developmental delay and the pupal lethality caused by loss of KDM5. Molecularly, our studies show that KDM5 functions by promoting the endoreplication of prothoracic gland cells, a process that increases ploidy and is rate-limiting for the expression of ecdysone biosynthetic genes. This occurs through KDM5-mediated regulation of the receptor tyrosine kinase *torso*, which in in turn promotes polyploidization and growth through activation of the MAPK signaling pathway. Taken together, our studies provide key insights into the biological processes regulated by KDM5 and the molecular mechanisms that govern the transcriptional regulation of animal development.

## INTRODUCTION

One key means by which transcription is regulated is through changes to covalent modifications that occur on nucleosomal histone proteins that comprise chromatin (Bannister and Kouzarides, 2011). The enzymes that add or remove these modifications play critical roles during development and their dysregulation can lead to disease states (Mirabella et al., 2016). Lysine demethylase 5 (KDM5) family proteins are chromatin-mediated regulators of transcription that are encoded by four paralogous genes in mammalian cells, KDM5A-D, and by a single gene in *Drosophila, kdm5* (also known as *little imaginal discs*; *lid*). The most well-established gene regulatory function of KDM5 proteins is their enzymatic activity that demethylates trimethylated lysine 4 of histone H3 (H3K4me3) (Accari and Fisher, 2015, Xhabija and Kidder, 2018). High levels of H3K4me3 are found surrounding transcriptional start sites and are associated with active gene expression (Santos-Rosa et al., 2002). Whereas absolute levels of H3K4me3 are unlikely to be primary drivers of gene expression levels, the breadth of these promoter peaks can impact transcriptional consistency (Benayoun et al., 2014, Howe et al., 2017). Recruitment of KDM5 to promoters to demethylate H3K4me3 is therefore one mechanism by which this family of proteins regulate transcription. KDM5 proteins can also affect gene expression through demethylase-independent mechanisms, such as by through interactions with the chromatin remodeling NuRD complex or by regulating histone acetylation via interactions with lysine deacetylase (HDAC) complexes (Barrett et al., 2007, Lee et al., 2009, Liu et al., 2014, Nishibuchi et al., 2014, Gajan et al., 2016).

KDM5 proteins play key roles in orchestrating diverse gene expression programs. This is emphasized by the large volume of literature linking dysregulation of KDM5 proteins to two seemingly disparate diseases, cancer and intellectual disability. Whole exome sequencing of patients with intellectual disability has identified loss of function mutations in KDM5A, KDM5B and KDM5C (Vallianatos and Iwase, 2015, Kim et al., 2017, Collins et al., 2019). Efforts to understand the link between KDM5 and intellectual disability using mice, flies and worms have revealed that KDM5 can regulate many genes that could impact neuronal development or function (Iwase et al., 2016, Zamurrad et al., 2018, Mariani et al., 2016, Chen et al., 2019). However, the extent to which any of these pathways contribute to the cognitive impairments observed in patients remains unknown. A similar deficit exists in our understanding of how the dysregulation of human KDM5 genes contributes to cancer (Blair et al., 2011, Plch et al., 2019). Unlike in intellectual disability, which is exclusively associated with loss of function mutations in KDM5 genes, malignancies have been associated with overexpression of KDM5A or KDM5B, either loss or gain of KDM5C, or loss of KDM5D. The best studied of these is the overexpression of KDM5B observed in breast cancer and melanoma that correlates with poor prognosis (Han et al., 2017). Despite being shown to directly or indirectly regulate genes involved in cell cycle progression, cancer stem cell survival and DNA repair, no clear model has emerged to explain its oncogenic capacities (Yamane et al., 2007, Catchpole et al., 2011, Roesch et al., 2013, Yamamoto et al., 2014, Han et al., 2017, McCann et al., 2019). Thus, our lack of understanding of KDM5-regulated pathways that lead to disease states underscores the importance of defining the physiological functions of KDM5 proteins.

*Drosophila melanogaster* offers a genetically amenable model to provide fundamental insight into KDM5 function *in vivo*, as it encodes a single, highly conserved, *kdm5* gene (Gildea et al., 2000). Moreover, in contrast to viable knockouts of KDM5A, KDM5B or KDM5C in mice, loss of *Drosophila* KDM5 results in lethality (Klose et al., 2007, Albert et al., 2013, Iwase et al., 2016, Drelon et al., 2018, Martin et al., 2018). This allows us to dissect critical functions of KDM5 without the complication of functional redundancy between mammalian KDM5 paralogs that could partially occlude phenotypes. We have previously shown that *kdm5* null mutants take 5 days longer than wild-type animals to complete larval development, linking KDM5 function to growth control. Interestingly, this phenotype is independent of the histone demethylase activity of KDM5, as animals specifically lacking this enzymatic function grow normally and produce viable adult flies (Li et al., 2010, Drelon et al., 2018). Emphasizing the importance of understanding cellular functions of KDM5 proteins that are independent of their enzymatic activity, the contribution of KDM5’ s demethylase function to normal development and to disease states in mammalian cells remains unresolved. For instance, while some intellectual disability-associated mutations in KDM5C reduce *in vitro* histone demethylase activity, others do not (Iwase et al., 2007, Tahiliani et al., 2007, Brookes et al., 2015, Vallianatos et al., 2018). Similarly, whereas the growth of some cancers can be attenuated by pharmacologically inhibiting KDM5 activity, KDM5 appears to act through demethylase-independent activities in others (Paroni et al., 2018, Cao et al., 2014). KDM5 proteins are therefore likely to utilize more than one gene-regulatory activity to control a range of cellular processes *in vivo.*

Here we demonstrate that KDM5 regulates larval development by playing an essential role in the prothoracic gland, a tissue that secretes the steroid hormone ecdysone, which is a key regulator of animal growth and maturation (Yamanaka et al., 2013). Although KDM5 is expressed ubiquitously during larval development, re-expression of *kdm5* exclusively in the prothoracic gland within a *kdm5* null mutant background rescues the larval growth delay and restores adult viability. We further show that within cells of the prothoracic gland, KDM5 is necessary to promote the endoreplicative cell cycles that increase DNA copy number and is required for the transcription of enzymes that mediate ecdysone production (Ohhara et al., 2017). This effect is likely to be achieved through regulation of the Torso receptor tyrosine kinase by KDM5. While stimulation of the Torso receptor and subsequent activation of the MAPK pathway is known to be necessary for ecdysone production (Rewitz et al., 2009), it has not been previously linked to either KDM5, nor to the regulation of prothoracic gland endocycles. Our data therefore provide new insight into the molecular mechanisms underlying the regulation of ecdysone production and developmental growth control.

## RESULTS

### KDM5 expression in the prothoracic gland is sufficient to rescue lethality and restore correct developmental timing to *kdm5* null mutants

To understand the underlying basis of the lethality caused by the *kdm5*^*140*^ null allele, we sought to define the spatial requirements of KDM5 function during development. To do this, we re-expressed *kdm5* in a null mutant background using a UAS-inducible transgene that we have previously used to rescue hypomorphic alleles of *kdm5* (Li et al., 2010) (Table 1). Ubiquitous expression of this UAS-*kdm5* transgene using Ubiquitin-Gal4 (Ubi-Gal4) resulted in slightly higher than endogenous levels of KDM5 expression and rescued the lethality of *kdm5*^*140*^ to produce morphologically normal adult flies (Figure 1A, B; Table 1). Re*-*expressing *kdm5* ubiquitously also led to a developmental timing profile that was indistinguishable from that of wild-type flies (Figure 1C). This is consistent with the observation by us and others that KDM5 is broadly expressed in all cell types examined to-date (Lee et al., 2009, Liu et al., 2014, Moshkin et al., 2009, Secombe et al., 2007, Tarayrah et al., 2015, Zamurrad et al., 2018).

**Table 1:** Re-expressing *kdm5* in defined tissues in *kdm5*^*140*^ mutants.

| Gal4 driver strain | Expression pattern of Gal4 driven expression of KDM5 | % of expected flies | Number of flies scored | Significant rescue of lethality? | Adult phenotype of rescued flies? |
| --- | --- | --- | --- | --- | --- |
| No Gal4 | None | 0 | 300 | - | - |
| Ubiquitin-Gal4 | Ubiquitous | 83.5% | 365 | P=8.8e-15 | Wild type |
| 32B-Gal4 | Wing imaginal disc<br>Salivary gland<br>Eye-antennal disc<br><u>Prothoracic gland</u> | 21.5% | 181 | P=6.8e-6 | Curved wings |
| T80-Gal4 | <u>Prothoracic gland</u><br>Imaginal discs<br>Muscle<br>Salivary gland<br>Larval brain<br>Gut | 77% | 132 | P=3.8e-15 | Wild type |
| Mai60-Gal4 | <u>Prothoracic gland</u><br>Larval brain<br>Imaginal disc | 62% | 246 | P=2.3e-15 | Curved wings |
| spok-Gal4 | <u>Prothoracic gland</u> | 39% | 173 | P=2.8e-9 | Curved wings |
| phm-Gal4 | <u>Prothoracic gland</u><br>Leg and wing discs<br>Trachea | 44.5% | 246 | P=1.4e-11 | Curved wings |
| CG-Gal4 | Fat body<br>Hemocytes<br>Lymph gland | 0% | 125 | ns P=1 | n.a. |
| dilp2-Gal4 | Insulin secreting cells of brain | 1.5% | 208 | ns P=1 | n.a. |
| Mef2-Gal4 | Somatic, visceral and cardiac muscle | 0% | 449 | P=1 | n.a. |
| D42-Gal4 | Motor neurons<br>Salivary gland | 0% | 222 | P=1 | n.a. |
| Aug21-Gal4 | Corpus allatum<br>Salivary gland<br>Tracheae<br>Malpighian tubules<br>Tv neurons | 2% | 173 | ns P=0.5 | n.a. |
| Akh-Gal4 | Corpus cardiaca | 0% | 139 | ns P=1 | n.a. |

Rescue of *kdm5*<sup>140</sup> lethality using the Gal4 drivers shown in the table shown as a percentage of the number of homozygous mutant *kdm5* flies expected based on Mendelian ratios. Significance determined using Fisher's exact test. n.a. not applicable.

**Figure 1:**
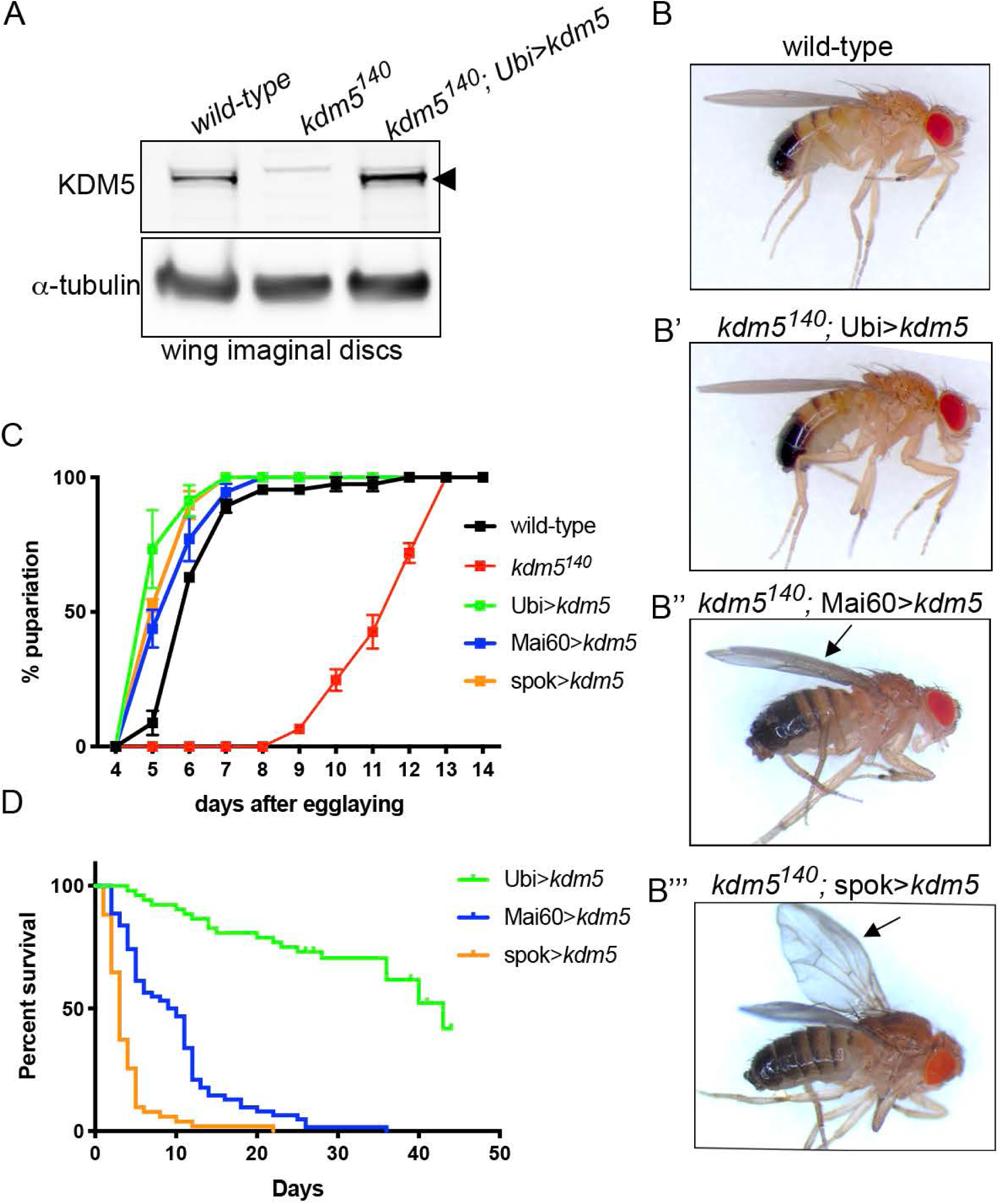
KDM5 expression in the prothoracic gland rescues *kdm5*^*140*^ developmental delay and lethality. (A) Western blot using 3^rd^ instar larval wing imaginal discs (five per lane) from wild-type, *kdm5*^*140*^ and *kdm5*^*140*^ expressing *kdm5* ubiquitously (*kdm5*^*140*^ ; Ubi>*kdm5*). Anti-KDM5 (top) is indicated by the arrowhead. Anti-alpha-tubulin is used as a loading control (bottom). (B) Male wild-type fly (*kdm5*^*140*^; g [*kdm5*: HA] attp86F). (B’) Male *kdm5*^*140*^, Ubi>*kdm5* adult fly showing morphologically normal features. (B’’) Male *kdm5*^*140*^; Mai60> *kdm5* adult fly with normal body and slightly curved wings indicated by arrow. (B’’’) Male *kdm5*^*140*^; spok> *kdm5* adult fly with normal body and curved wings indicated by arrow. (C) Number of days taken for pupariation to occur in wild type (N=82), *kdm5*^*140*^ (N=49), *kdm5*^*140*^ ; Ubi>*kdm5* (N=69), *kdm5*^*140*^ ; Mai60>*kdm5* (N=63) and *kdm5*^*140*^ ; spok>*kdm5* (N=74). (D) Adult survival for *kdm5*^*140*^; Ubi> *kdm5* (N=56), *kdm5*^*140*^; Mai60>*kdm5* (N=62) and *kdm5*^*140*^; spok> *kdm5* (N=51) rescued male flies. Mai60 and spok-Gal4-rescued flies are significantly shorter lived than Ubi-Gal4-rescued flies (Mantel-Cox log-rank test; P< 0.0001).

To test whether KDM5 plays key developmental roles in specific tissues, we tested a range of Gal4 drivers for their ability to rescue *kdm5*^*140*^ lethality when combined with UAS-*kdm5* (Table 1). These included strains that drive Gal4 expression in tissues that have phenotypes in *kdm5* mutant or knockdown larvae such as imaginal discs and hemocytes (Drelon et al., 2018, Moran et al., 2015, Moshkin et al., 2009). In addition, we tested tissues linked to the regulation of larval growth such as the hormone-producing larval prothoracic gland, insulin secreting cells of the brain, or cells of the fat body that coordinate larval growth with feeding and nutritional status (Yamanaka et al., 2013). Significantly, all Gal4 drivers that were expressed in the larval prothoracic gland significantly rescued *kdm5*^*140*^ lethality, including spookier-Gal4 (spok-Gal4), which is expressed exclusively in this tissue (Table 1) (Hrdlicka et al., 2002, Moeller et al., 2017, Shimell et al., 2018). Consistent with KDM5 playing critical functions in the ecdysone-secreting larval prothoracic gland, spok-Gal4-mediated re-expression of *kdm5* was sufficient to rescue the developmental delay of *kdm5*^*140*^ (Figure 1C). It is, however, notable that while expression of *kdm5* in the prothoracic gland allowed *kdm5*^*140*^ mutant pupae to eclose, adult flies were shorter-lived than flies expressing *kdm5* ubiquitously (Figure 1D). In addition, while adults rescued by ubiquitous expression of *kdm5* were largely male and female fertile, prothoracic gland re-expression of *kdm5* resulted in significant infertility (Table 2). These data are consistent with previous observations showing roles for KDM5 in oogenesis and in testis germline stem cell proliferation (Tarayrah et al., 2015, Navarro-Costa et al., 2016, Zhaunova et al., 2016). Together, these studies demonstrate essential roles for KDM5 in the larval prothoracic gland and in other cell types that are important for wing maturation, adult survival and reproduction.

**Table 2:** Fertility of *kdm5*^*140*^-rescued adult flies.

| Gal4 driver strain | % of fertile female flies | % of fertile male flies |
| --- | --- | --- |
| **Ubi-Gal4** | 82% (N=17) | 79% (N=19) |
| **Mai60-Gal4** | 7% (N=14) P=3.8e-05 | 0% (N=14) P=6e-06 |
| **spok-Gal4** | 0% (N=9) P=7e-05 | 0% (N=17) P=7.2e-07 |

Fertility of *kdm5*<sup>140</sup> homozygous mutant male and female flies expressing UAS-*kdm5* under the control of Ubi-Gal4, spok-Gal4 or Mai60-Gal4. P-values were calculated using Fisher's exact test compared to Ubi-Gal4-rescued control flies.

These rescue studies also revealed a role for KDM5 in wing development that is not essential for viability. While adult flies generated by re-expression of *kdm5* in the prothoracic gland such as spok-Gal4 were the same size as those rescued by ubiquitous expression, they had wings that were curved downward (Figure 1B). A similar curved-down wing phenotype was observed in *kdm5*^*140*^ flies that expressed *kdm5* in the wing imaginal disc in addition to the prothoracic gland such as Mai60-Gal4 or 32B-Gal4, but not with more broadly expressed drivers like T80-Gal4 (Figure 1B, Table 1). This curved-wing phenotype could be caused by the proliferative or cell death phenotypes we previously observed in *kdm5*^*140*^ larval wing imaginal discs (Drelon et al., 2018). To test this, we quantified wing disc proliferation and cell death using antibodies that detect the mitotic marker phosphorylated histone H3 (pH3) and the cleaved caspase Dcp-1, respectively. As expected, the decreased levels of pH3 observed in *kdm5*^*140*^ mutant wing imaginal discs were fully restored by ubiquitous *kdm5* expression using Ubi-Gal4 (Figure 2A-C). Significantly, expressing *kdm5* under the control of Mai60-Gal4 or spok-Gal4, which drive low or no KDM5 expression in the wing disc, also rescued pH3 to wild-type levels (Figure 2B, C). Similarly, the increased number of Dcp-1 positive cells seen in *kdm5*^*140*^ wing discs was restored to wild type levels by Mai60-Gal4 or spok-Gal4 driven expression of *kdm5* (Figure 2D-I). Thus, neither the proliferative nor the apoptotic phenotypes of *kdm5*^*140*^ wing imaginal discs are triggered by loss of KDM5 in the disc itself but are caused by cell-nonautonomous mechanisms. While the basis for the wing defects observed in Mai-60 and spok-Gal4 rescued flies remains unclear, it could be due to a requirement for KDM5 during pupal development or during wing maturation in newly eclosed flies.

**Figure 2:**
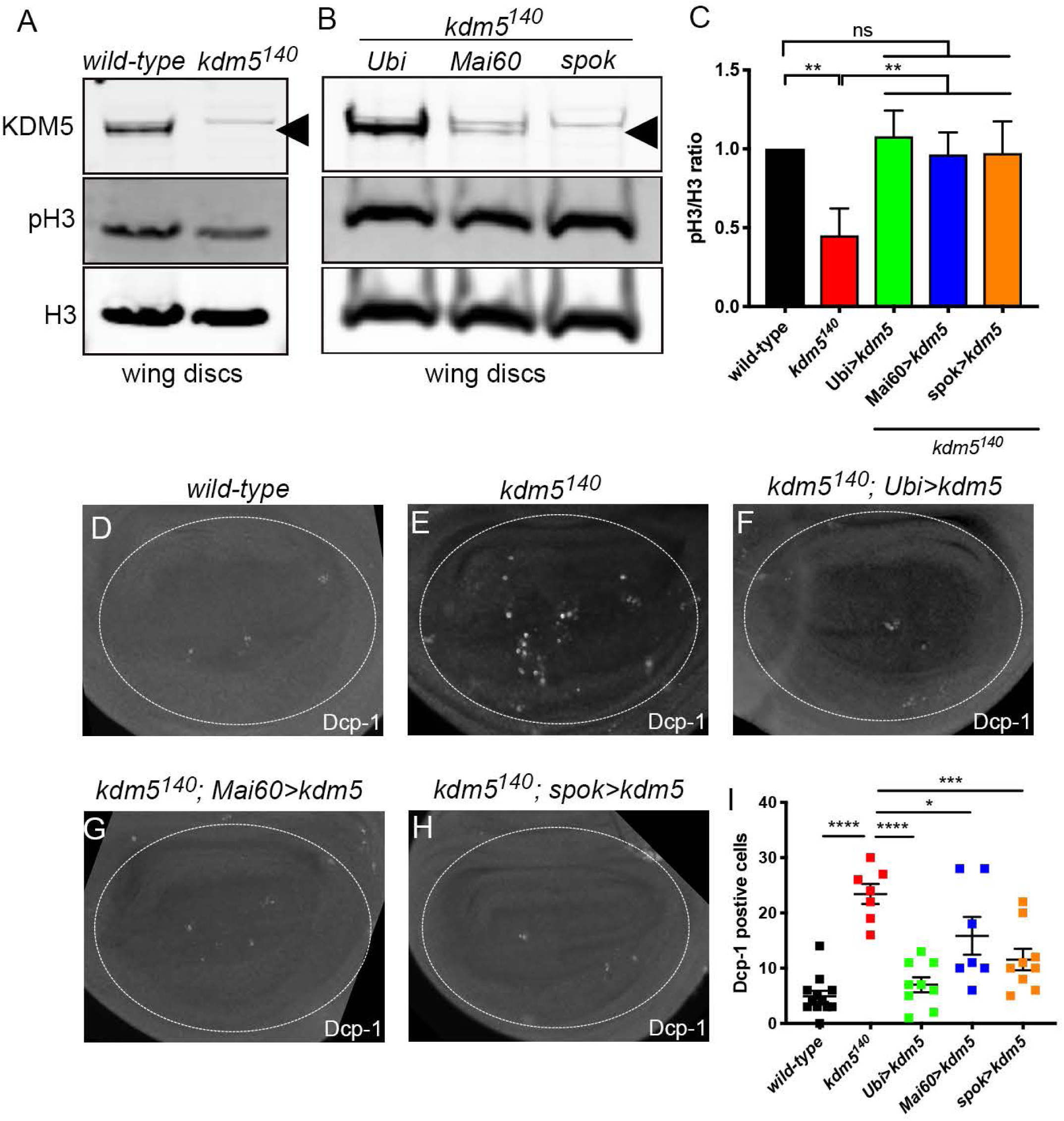
Increased cell death and reduced proliferation in *kdm5*^*140*^ wing discs is due to non-cell autonomous effects. (A) Western blot using anti-KDM5 (top), anti-phosphorylated histone H3 (middle; pH3) and total histone H3 (bottom; H3) in wild-type and *kdm5*^*140*^ wing discs. Six wing discs per lane. (B) Western blot using wing discs from *kdm5*^*140*^ mutants re-expressing *kdm5* using Ubi-Gal4, Mai60-Gal4 or spok-Gal4. anti-KDM5 (top), pH3 (middle) and total H3 (bottom). Six wing discs per lane. (C) Quantification of the levels of pH3 relative to total H3 from three Western blots. * * P <0.01 (one-way ANOVA). Error bars indicate SEM. (D) Wild-type wing imaginal disc from wandering 3^rd^ instar larva stained with anti-Dcp-1. Dotted circle indicates pouch region of the wing disc that was used to count Dcp-1 positive cells. (E) *kdm5*^*140*^ mutant wing imaginal disc stained with the apoptosis marker anti-Dcp-1. (F) Anti-Dcp-1 staining of a wing imaginal disc from *kdm5*^*140*^ mutant re-expressing *kdm5* in imaginal discs and prothoracic gland using Mai60-Gal4 (*kdm5*^*140*^; Mai60> *kdm5*). (G) Anti-Dcp-1 staining of a wing imaginal disc from *kdm5*^*140*^ mutant re-expressing *kdm5* in the prothoracic gland with spok-Gal4 (*kdm5*^*140*^ ; spok> *kdm5*). (H) Quantification of the number of Dcp-1 positive cells in the pouch region of wing imaginal discs from wild type (N=12), *kdm5*^*140*^ (N=7), *kdm5*^*140*^ ; Ubi>*kdm5* (N=9), *kdm5*^*140*^ ; Mai60>*kdm5* (N=7) and *kdm5*^*140*^ ; spok>*kdm5* (N=8). ****P<0.0001, ***P<0.001, *P<0.05 (one-way ANOVA). Error bars indicate SEM.

### Reduced ecdysone production causes the developmental delay of *kdm5*^*140*^ mutant larvae

The prothoracic gland, corpus allatum and corpora cardiaca are the three sub-tissues that comprise the larval ring gland, a semi-circular structure associated with the central larval brain. Whereas the prothoracic gland synthesizes ecdysone, the corpus allatum and corpora cardiaca produce juvenile hormone and adipokinetic hormone, respectively (Yamanaka et al., 2013). In order to evaluate the role of KDM5 in these two other sub-tissues, we tested whether expressing *kdm5* in the corpus allatum rescued the lethality of *kdm5*^*140*^ using *Aug21*-Gal4 (Table 1) (Gaziova et al., 2004). In contrast to expression in the prothoracic gland, no significant rescue was observed when *kdm5* was reintroduced in the corpus allatum. Similarly, expression of *kdm5* in the corpora cardiaca using Akh-Gal4 failed to rescue *kdm5*^*140*^ mutants (Lee and Park, 2004) (Table 1). Thus, within the ring gland, KDM5 is critical only for prothoracic gland function, linking its transcriptional regulatory functions to ecdysone biology.

Based on the requirement for KDM5 in the prothoracic gland, we tested whether *kdm5*^*140*^ larvae had altered levels of the active form of ecdysone (20-hydroxyecdysone; 20E). These analyses revealed that *kdm5*^*140*^ larvae had a 3-fold reduction in 20E levels and concomitant reduction in the expression of ecdysone-regulated genes such as *broad, E74* and *E75* (Figure 3A, B). Ecdysone is synthesized in the prothoracic gland from cholesterol through the action of a series of cytochrome p450 biosynthetic enzymes that include *neverland* (*nvd*), *spookier* (*spok*) and *disembodied* (*dib*) (Gilbert, 2004). It is then subsequently converted to 20E in cells of the larva by the hydroxylase Shade (*shd*). qPCR analyses to examine the expression of these genes revealed that *nvd, spok* and *dib* transcripts were significantly decreased in *kdm5*^*140*^ mutant 3^rd^ instar larvae, whereas levels of *shd* mRNA were unchanged (Figure 3C, D). *kdm5*^*140*^ mutants therefore have reduced 20E due to a defect in the biosynthesis of ecdysone in the prothoracic gland.

**Figure 3:**
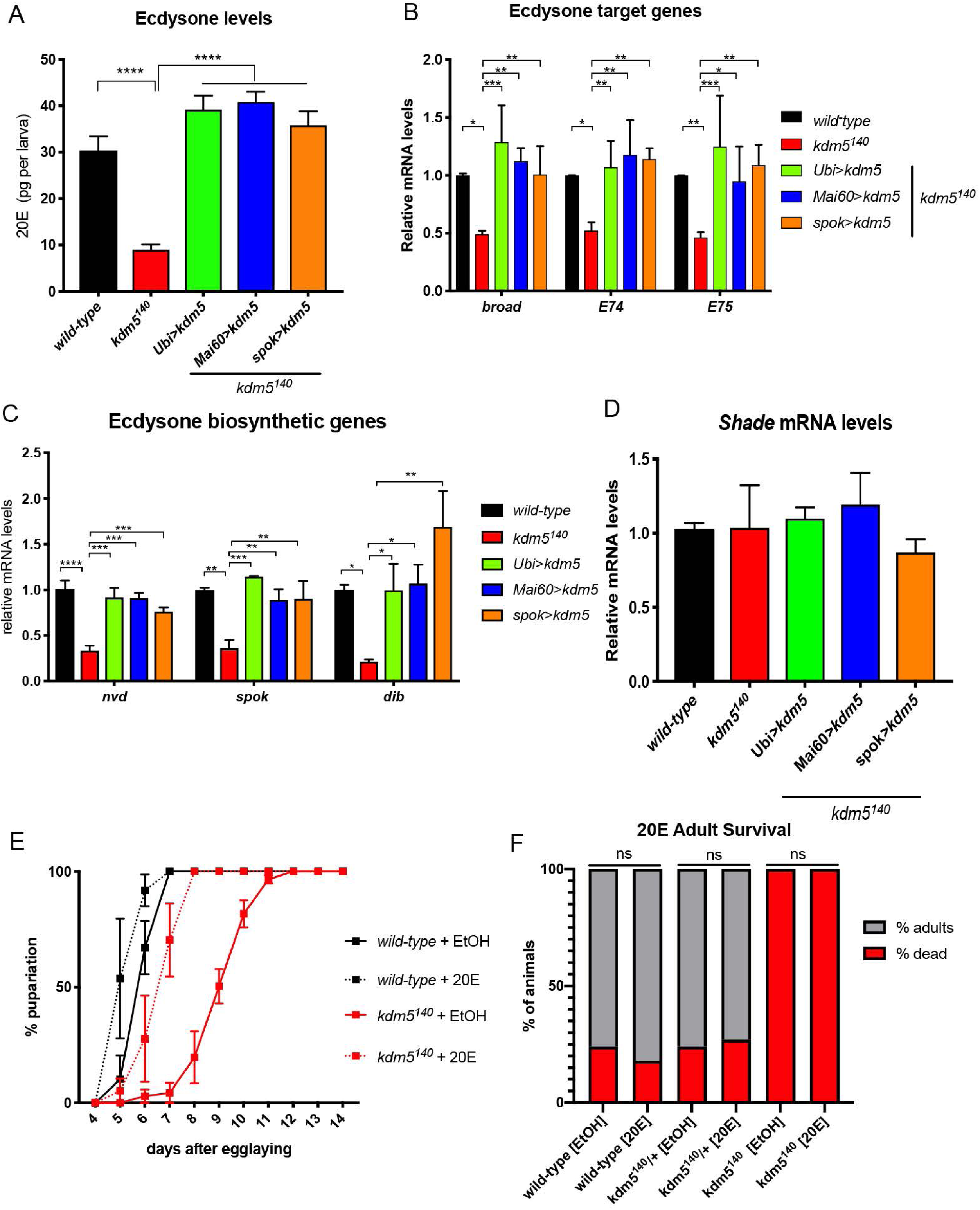
Exogenous ecdysone rescues the developmental delay of *kdm5*^*140*^ mutants. (A) Quantification 20E levels in pg per larva using wild-type, *kdm5*^*140*^, *kdm5*^*140*^; Ubi> *kdm5, kdm5*^*140*^; Mai60> *kdm5* and *kdm5*^*140*^; spok> *kdm5* whole 3^rd^ instar larvae. * * * * P< 0.001 (one-way ANOVA). Error bars indicate SEM. (B) Real-time PCR from biological triplicate samples quantifying the mRNA levels of the ecdysone target genes *broad, E74* and *E75* in wild type, *kdm5*^*140*^, *kdm5*^*140*^ ; Ubi>*kdm5, kdm5*^*140*^ ; Ma60>*kdm5* and *kdm5*^*140*^ ; spok>*kdm5* whole larvae. Data were normalized to *rp49* and shown relative to wild type. * P<0.05, **P<0.01, ***P<0.001. Error bars indicate SEM. (C) Real-time PCR from biological triplicate showing mRNA levels *nvd, spok* and *dib* from whole 3^rd^ instar wild-type, *kdm5*^*140*^ ; Ubi>*kdm5, kdm5*^*140*^ ; Mai60>*kdm5* and *kdm5*^*140*^ ; spok>*kdm5* larvae. Data were normalized to *rp49* and shown relative to wild type. *P=0.05, **P<0.01, ***P<0.001, **** P<0.0001. Error bars indicate SEM. (D) Real-time PCR from biological triplicate showing mRNA levels of the hydroxylase *Shade* from whole 3^rd^ instar wild type, *kdm5*^*140*^ ; Ubi>*kdm5, kdm5*^*140*^ ; Mai60>*kdm5* and *kdm5*^*140*^ ; spok>*kdm5* larvae. Data were normalized to *rp49* and shown relative to wild type. Error bars indicate SEM. (E) Quantification of the time for pupariation to occur upon feeding 20E to wild type (N=103) and *kdm5*^*140*^ (N=62) or vehicle alone (ethanol; EtOH) to wild type (N=67) or *kdm5*^*140*^ (N=60). (F) Quantification of the number of pupae that eclose from their pupal case for wild-type larvae that were fed EtOH vehicle (N=33) or 20E (N=40), *kdm5*^*140*^ heterozygotes fed EtOH (N=165) or 20E (N=98) and *kdm5*^*140*^ homozygotes from the same vial as heterozygotes fed EtOH (N=16) or 20E (N=15). ns, not significant (Fisher’ s exact test).

To test whether re-expression of *kdm5* in the prothoracic gland restored ecdysone production, we quantified levels of 20E in *kdm5*^*140*^ larvae expressing UAS-*kdm5* using Ubi-Gal4, Mai60-Gal4 or spok-Gal4. All three Gal4 drivers restored levels of 20E, the expression of ecdysone-regulated genes, and transcription of the ecdysone biosynthetic genes *nvd, spok* and *dib* to wild type levels (Figure 3A-D). Based on these data, dietary supplementation with 20E would be expected to attenuate the phenotypes caused by loss of KDM5. We therefore fed wild-type and *kdm5*^*140*^ larvae food containing 20E. As shown in Figure 3E, exogenously providing 20E rescued the developmental delay of *kdm5* mutant larvae compared to a vehicle alone control, emphasizing the importance of KDM5-mediated regulation of ecdysone production. 20E was not, however, able to rescue the lethality of *kdm5*^*140*^ animals whereas wild-type and *kdm5* heterozygous animals eclosed after ecdysone feeding (Figure 3F). Loss of KDM5 therefore causes additional defects in the prothoracic gland that are not able to be substituted by feeding 20E.

### KDM5 regulates the Torso/MAPK signaling pathway and prothoracic gland endocycles

To investigate the basis of the reduced ecdysone levels, we examined the activity of two key pathways known to regulate its production: the Torso/MAPK and insulin receptor pathways (Yamanaka et al., 2013). Activation of the insulin receptor by insulin-like peptides leads to the phosphorylation of Akt, which subsequently induces cell growth and ecdysone production (Ohhara et al., 2017, Danielsen et al., 2016). To test whether this pathway was affected by the loss of KDM5, we examined levels of phosphorylated Akt compared to total Akt in *kdm5*^*140*^ ring glands by Western blot (Figure 4A, B). We also quantified mRNA levels of the insulin receptor (InR) in brain-ring gland complexes (Figure 4C). Neither of these signaling components were affected in *kdm5*^*140*^, suggesting that this pathway was unlikely to mediate the developmental delay caused by loss of KDM5.

**Figure 4:**
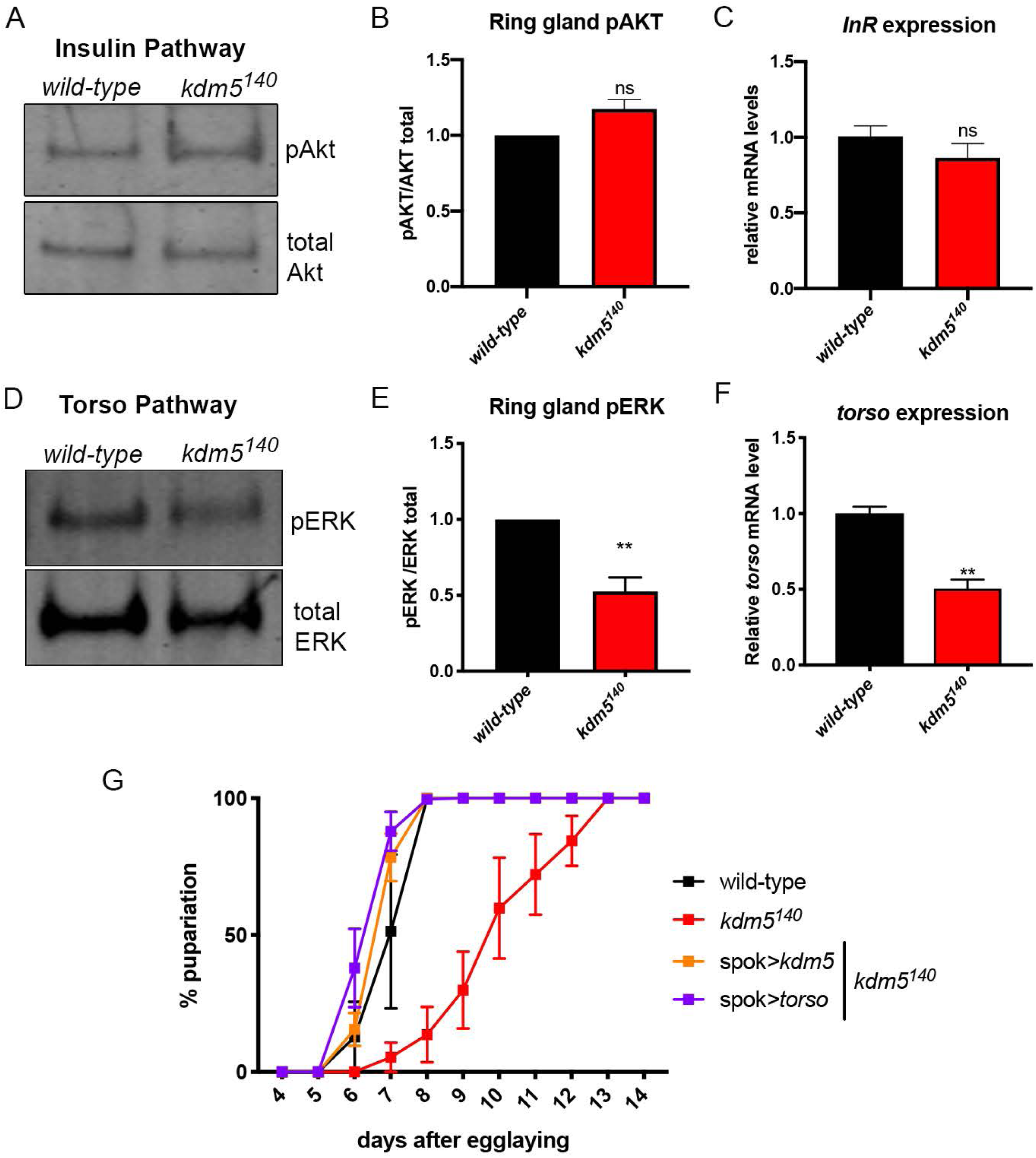
Levels and activity of the Torso receptor pathway are reduced in *kdm5*^*140*^. (A) Western blot from wild-type and *kdm5*^*140*^ ring glands showing levels of phosphorylated Akt and total Akt as a control. Six rings glands per lane. (B) Quantitation of triplicate Western blots showing ratio of pAkt to total Akt in *kdm5*^*140*^ and wild type. ns not significant. (C) Real-time PCR quantitation showing average of biological triplicate of *InR* mRNA levels from dissected brain-ring gland complexes from wild-type and *kdm5*^*140*^ larvae. Data were normalized to *rp49* and shown relative to wild type. ns not significant. Error bars indicate SEM. (D) Western blot analyses of wild-type and *kdm5*^*140*^ dissected ring glands examining levels of phosphorylated ERK (pERK; top) and total ERK (bottom). Five ring glands per lane. (E) Quantitation of the ratio of pERK to total ERK in *kdm5*^*140*^ compared to wild-type from three Western blots. * * P< 0.01 (Student’ s t-test). Error bars indicate SEM. (F) Real-time PCR quantitation showing average of biological triplicate of *torso* mRNA levels from dissected brain-ring gland complexes from wild-type and *kdm5*^*140*^. Data were normalized to *rp49* and shown relative to wild type. * * P=0.0026 (Student’ s t-test). Error bars indicate SEM. (G) Number of days taken for pupariation to occur for wild-type (N=87) *kdm5*^*140*^ (N=46), *kdm5*^*140*^; spok> *kdm5* (N=57), and *kdm5*^*140*^; spok> *torso* (N=73) animals.

Ecdysone production can also be regulated by activation of the Torso receptor by prothoracicotropic hormone (PTTH), which induces a Ras/ERK MAP kinase cascade (Niwa and Niwa, 2014, Yamanaka et al., 2013). To test whether this pathway was altered in *kdm5* mutant larvae, we examined levels of phosphorylated ERK relative to total ERK in addition to mRNA levels of the *torso* receptor (Figure 4D, E, F). Both phosphorylated ERK and *torso* expression were significantly decreased compared to wild type, suggesting that loss of KDM5 caused a defect in this pathway. To examine the functional role that the Torso/MAPK pathway plays in the developmental delay and lethality of *kdm5*^*140*^, we expressed a UAS transgene encoding wild-type *torso* or an activated form of the downstream GTPase Ras (Ras^V12^). Expression of either of these transgenes restored the larval growth defect and the lethality of *kdm5*^*140*^ and produced adults that, like reintroduction of *kdm5*, showed a curved-down phenotype (Figure 4G; Table 3). These data are consistent with the reduced Torso/MAPK signaling observed in *kdm5*^*140*^ playing a key role in mediating KDM5’ s prothoracic gland activities.

**Table 3:** Rescue of *kdm5* lethality by UAS-transgene expression.

| UAS-transgene expressed under the control of spok-Gal4 | % of expected flies | Number of flies scored | Significant rescue of lethality? | Adult phenotype of rescued flies? |
| --- | --- | --- | --- | --- |
| UAS-*torso* | 13% | 299 | P=0.002 | Curved wings |
| UAS-*Ras*<sup>V12</sup> | 31% | 90 | P=9.2e-8 | Curved wings |
| UAS-*cyclin E* | 0% | 200 | ns P=1 | n.a. |

Rescue of *kdm5*<sup>140</sup> lethality by expressing the UAS transgene indicated. Data are shown as a percentage of the number of homozygous mutant flies expected based on Mendelian genetics. Fisher's exact test was used to determine whether the rescue was significant compared to *kdm5*<sup>140</sup> flies harboring no UAS transgene. n.a. not applicable.

While Torso/MAPK activation is known to lead to ecdysone production (Rewitz et al., 2009), the mechanisms underlying this link are unclear. One requirement for maximal expression of the genes that are limiting for the synthesize ecdysone are a series of endoreplicative cell cycles that increases the DNA content of prothoracic cells up to ∼64C (Ohhara et al., 2017, Aggarwal and King, 1969). Loss of KDM5 could therefore affect ecdysone production by affecting the ability of prothoracic gland cells to endoreplicate, and ectopic activation of the Torso/MAPK pathway may be able to restore this defect. To test whether KDM5 was required for polyploidization, we quantified the number of *kdm5* mutant prothoracic gland cells undergoing S-phase by incorporating the thymine nucleoside analog 5-ethynyl-2’-deoxyuridine (EdU). This revealed a significant decrease in the number of EdU positive cells in *kdm5*^*140*^ compared to wild type (Figure 5A-I). Importantly, loss of KDM5 caused these endocycle defects without altering the total number of cells that comprise the prothoracic gland (Figure 5J). The specification of these cells therefore occurs normally during embryogenesis in *kdm5* mutants, but fail to endocycle and grow correctly during larval development. We next tested the extent to which *torso* expression in the prothoracic gland rescued the endocycling defect of *kdm5*^*140*^. Consistent with the notion that the Torso/MAPK pathway affects ecdysone by promoting endoreduplication, expression of *torso* using spok-Gal4 was able to restore prothoracic gland polyploidization in a similar manner to expression of *kdm5* (Figure 5I).

**Figure 5:**
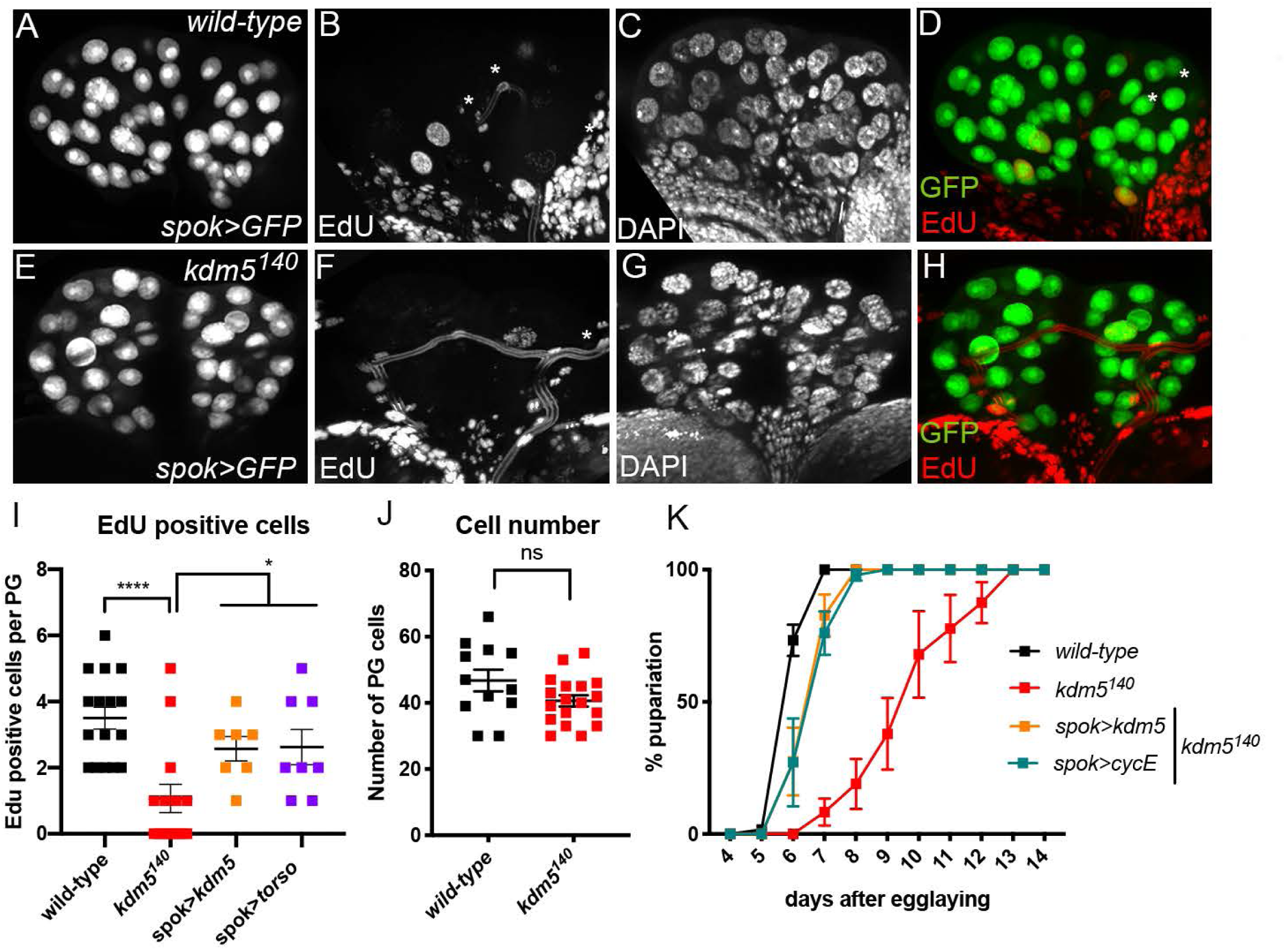
*kdm5*^*140*^ larvae have a prothoracic gland endocycle defect that is rescued by expression of *cyclin E* or *torso*. (A-D) Control flies in which EdU incorporation was carried out using larvae from a cross of spok>GFP to the wild-type strain *w*^*1118*^ to mark prothoracic gland cells. (A) GFP (B) EdU (C) DAPI (D) merge of EdU and GFP channels. (E-H) EdU incorporation into *kdm5*^*140*^ mutant larva carrying spok>GFP to mark cells of the prothoracic gland. (E) GFP (G) EdU (G) DAPI (H) merge of EdU and GFP channels. (I) Quantification of the number of EdU positive prothoracic gland nuclei from control larvae (N=7 larvae) and *kdm5*^*140*^ homozygous mutant larvae (N=10), *kdm5*^*140*^; spok*> kdm5* (N=7) and *kdm5*^*140*^; spok*> torso* (N=8). In all genotypes, prothoracic gland cells were marked using spok> GFP. * * * * P< 0.0001, * P< 0.05 (one-way ANOVA). Error bars indicate SEM. (J) Quantification of the total prothoracic gland cell number in control spok> GFP (N=12 larvae) and *kdm5*^*140*^; spok> GFP (N=18) larvae. Error bars indicate SEM. ns not significant (Student’ s t-test). (K) Quantification of the time for pupariation to occur in wild-type (N=91), *kdm5*^*140*^ (N=56), *kdm5*^*140*^; spok> *kdm5* (N=73) and *kdm5*^*140*^; spok> *cyclin E* (N=48) animals.

To further explore the link between phenotypes caused by loss of KDM5 and reduced prothoracic gland endocycling, we directly drove endoreplication by overexpressing the cell cycle regulator Cyclin E (Ohhara et al., 2017). Spok-Gal4-mediated expression of a UAS-*cyclin E* transgene in *kdm5*^*140*^ mutant larvae was sufficient to rescue their slowed larval development (Figure 5K). A deficit in prothoracic gland cell endocycling is therefore likely to be a key cause of the developmental delay seen in *kdm5*^*140*^ larvae. Significantly, in contrast to activation of the Torso/MAPK pathway, expression of *cyclin E* was not able to restore viability to *kdm5*^*140*^ mutant animals (Table 3). KDM5 therefore likely regulates larval growth rate by activating the *torso*/MAPK pathway, which promotes the polyploidization of prothoracic gland cells that is limiting for the production of steroidogenic enzymes (Figure 6). These data also indicate that *kdm5* mutant larvae have currently unknown prothoracic gland defects that are not able to be overcome by restoring ecdysone levels, but can be rescued by activating the MAPK pathway by overexpressing *torso* or *Ras*^*V12*^.

**Figure 6:**
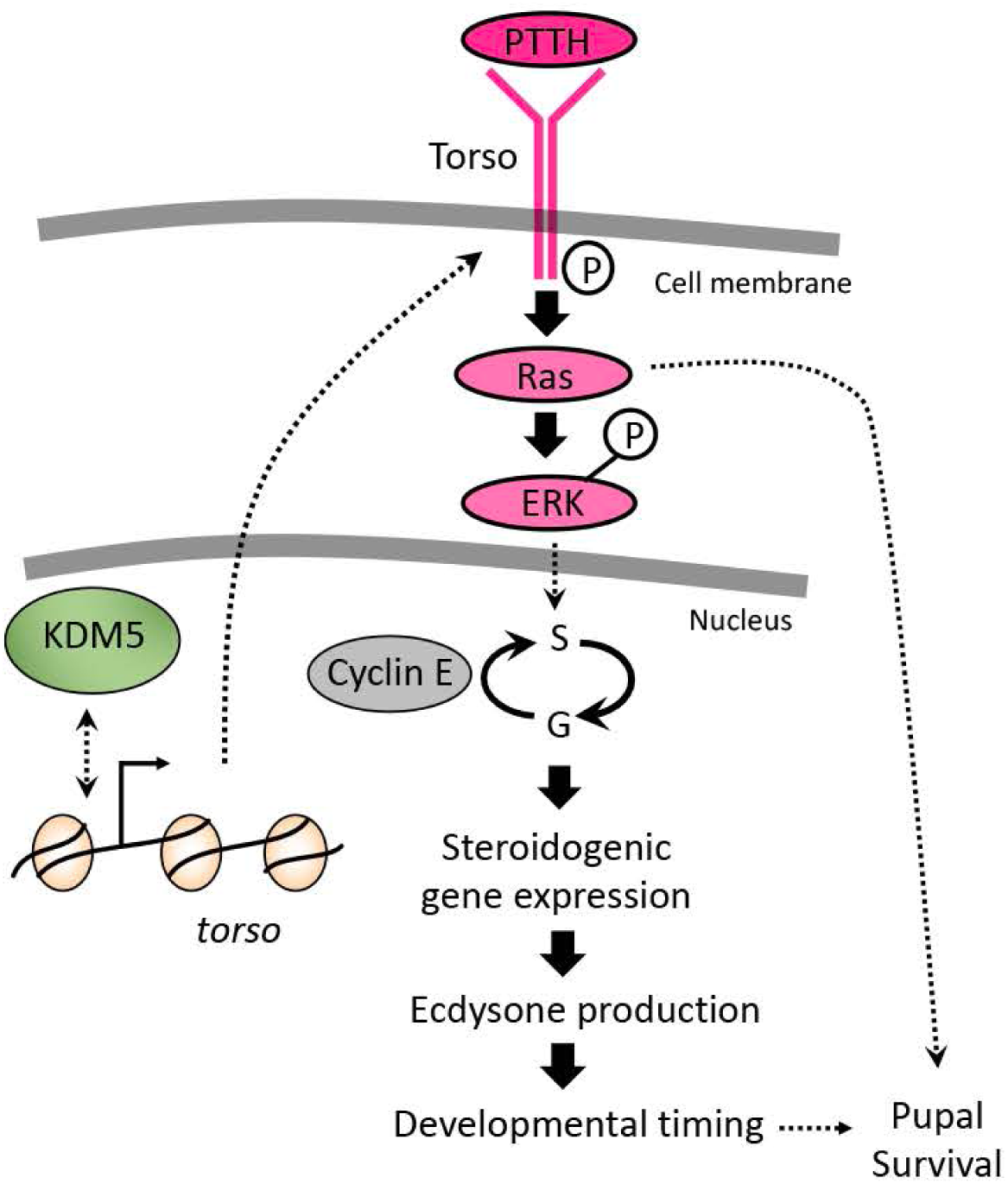
Model for KDM5 function in the prothoracic gland. We propose that KDM5 activates the Torso/MAPK pathway to regulate prothoracic gland endocycling and subsequent ecdysone production in addition to other key cellular functions.

## Discussion

In this study, we demonstrate that KDM5 is essential for the function of the ecdysone-producing prothoracic gland during *Drosophila* larval development. Critical to this conclusion was our finding that expressing *kdm5* in the prothoracic gland was sufficient to rescue the lethality and developmental delay phenotypes of *kdm5*^*140*^ null allele homozygous mutant animals. Consistent with this observation, prothoracic gland function was defective in *kdm5* mutants, with mutant larvae having low levels of ecdysone and reduced expression of downstream hormone-responsive target genes. Demonstrating the importance of KDM5-mediated regulation of ecdysone production, dietary supplementation of 20E restored normal developmental timing to *kdm5* mutant larvae. 20E was not, however, able to restore adult viability to *kdm5* mutant animals. At the cellular level, loss of KDM5 slowed the prothoracic gland endoreplicative cycles that increase the ploidy of these cells and are key to ecdysone biosynthesis (Ohhara et al., 2017). Restoring these endocycles reestablished normal developmental timing to *kdm5*^*140*^ mutants but was not able to rescue their lethality. In contrast, ectopic activation of the Torso/MAPK pathway was able to restore both developmental timing and rescue the lethality caused by loss of KDM5. We therefore propose that KDM5-mediated activation of the Torso pathway is essential for larval growth control by promoting prothoracic gland polyploidization and that KDM5 also affects additional cellular process(es) that are critical to survival.

We suggest that one key function of KDM5 is to regulate the expression of the *torso* gene, which then affects downstream MAPK signaling. Consistent with this view, mutations in *torso*, or ablation of the neurons that produce its ligand PTTH, cause a 5-day delay to larval development similar to that observed for *kdm5*^*140*^ (McBrayer et al., 2007, Drelon et al., 2018, Rewitz et al., 2009). Little is known about the transcriptional regulation of *torso* and other genes that comprise the pathways that mediate ecdysone production. One possibility is that KDM5 is a direct transcriptional activator of the *torso* gene, thus decreased expression of this receptor would be expected in *kdm5* mutant animals. Because of the small size of the prothoracic gland, it is not currently feasible to carry out ChIP experiments to examine KDM5 promoter binding in this tissue. It is, however, notable that the promoter of *torso* was not bound by KDM5 in existing ChIP-seq datasets from the larval wing imaginal discs or from whole adult flies (Liu and Secombe, 2015, Lloret-Llinares et al., 2012). This could be because *torso* expression is largely restricted to the prothoracic gland during larval development and so may not be expected to have promoter-bound KDM5 in the tissues examined to-date (Rewitz et al., 2009). Alternatively, KDM5 might regulate *torso* indirectly. In the silkworm *Bombyx mori*, expression of the *torso* gene is repressed in response to starvation conditions (Gu et al., 2011). Although the mechanism by which this occurs is unknown, it does indicate that other cellular defects caused by loss of KDM5 could lead to changes to *torso* transcription and subsequent decrease in ecdysone production. Whether the regulation of *torso* by KDM5 is direct or indirect, it occurs in a demethylase-independent manner, as larvae lacking enzymatic activity show a normal developmental profile (Drelon et al., 2018). Consistent with this observation, components of KDM5 complexes that regulate gene expression through demethylase-independent mechanisms also affect developmental timing. For example, a development delay similar to that of *kdm5*^*140*^ is caused by RNAi-mediated knockdown of the histone deacetylase *HDAC1* or the NuRD complex components *asf1* and *Mi-2* (Danielsen et al., 2016). KDM5 could therefore interact with these proteins to regulate the expression of genes critical to the regulation of larval development.

Our data show that KDM5-mediated regulation of the Torso/MAPK pathway affects ecdysone production by promoting prothoracic gland endoreplication. While Torso/MAPK has been proposed to promote ecdysone production by activating the expression of ecdysone biosynthetic genes (Rewitz et al., 2009, Danielsen et al., 2013), it has not been linked to the regulation of polyploidization. Our model is, however, consistent with recent data demonstrating that increased ploidy within the cells of the prothoracic gland is necessary for the optimal expression of steroidogenic genes (Ohhara et al., 2017). Because the genes required for ecdysone biosynthesis are among the most abundantly expressed in the prothoracic gland, their mRNA levels may be entirely limited by gene copy number (Christesen et al., 2017, Fox and Duronio, 2013). A similar requirement for copy number amplification to produce peak gene expression levels has been observed in other cell types in *Drosophila*, including chorion gene expression in ovarian follicle cells (Orr-Weaver, 2015). How could Torso/MAPK activation promote prothoracic gland cell cycle progression? One possibility is by affecting levels of cell cycle regulators such as the transcription factor E2f1, which is essential for both mitotic and endoreplicative cell cycles (Davoli and de Lange, 2011, Fox and Duronio, 2013, Frawley and Orr-Weaver, 2015). This model is based on studies of the polyploid enterocytes of the adult midgut, in which activation of the MAPK pathway via the EGF receptor stabilizes E2f1 protein, leading to transcription activation of *cyclin E* (Xiang et al., 2017). Because restoring the polyploidization defect of *kdm5* mutant prothoracic glands restored developmental timing but not lethality, this model would require that ectopic Torso/MAPK also impact the activity of additional cellular pathways in concert with, or independent from, a direct effect on cell cycle regulators. A potential candidate for this is the target of rapamycin (TOR) pathway, which serves as a hub for the integration of extrinsic and intrinsic signals in many cell types, including the prothoracic gland (Saxton and Sabatini, 2017b). The TORC1 complex is required for expression of *cyclin E*-mediated prothoracic gland polyploidization (Ohhara et al., 2017) and is also linked to numerous downstream cellular consequences including autophagy, changes to translation and to cell growth and division (Saxton and Sabatini, 2017a). An effect of KDM5 on TOR activity would provide a means by which KDM5 affects numerous key cellular processes. Genetic dissection of where KDM5 lies with respect to established components of the Torso and TOR pathways will provide clarity on this important issue.

Our observed role of KDM5 in the growth and polyploidization of larval prothoracic gland cells raises the possibility that it might play a role in other cell types that utilize endoreplicative cycles. This could have broad consequences for our understanding of KDM5 biology, since polyploidization is observed in many plant and animal cell types and is widely used during *Drosophila* larval development (Orr-Weaver, 2015, Fox and Duronio, 2013). For example, polyploid cells of the larval fat body survive until early adulthood where they are necessary for survival because flies are unable to immediately begin feeding (Aguila et al., 2007). In fact, a role for KDM5 in cells of the fat body would provide an explanation for the shortened adult lifespan of *kdm5* mutant flies rescued by prothoracic gland-specific expression of *kdm5*. In addition to being critical for normal development, endocycling cells are also found in tumors in both humans and *Drosophila* (Fox and Duronio, 2013, Davoli and de Lange, 2011, Cong et al., 2018). While the role of polyploid cells in the etiology or maintenance of cancers remains a topic of ongoing research, KDM5-regulated endocycling could contribute to its tumorigenic activities in humans. Regulation of polyploidization the nervous system could also contribute to the role of KDM5 proteins in intellectual disability (Vallianatos and Iwase, 2015). This could, for instance, be mediated by KDM5 function in glial cells, as polyploidization of a superineurial glial cells in *Drosophila* is required for normal brain development (Unhavaithaya and Orr-Weaver, 2012). While it is not clear the extent to which a similar phenomenon occurs during human brain development, it is interesting to note that glial cell types contribute to the clinical severity of intellectual disability disorders such as Rett syndrome (Sharma et al., 2018). Thus, while there is still much to be learned regarding the contribution of polyploid cells to normal development and to disease states, KDM5-regulated transcriptional programs are likely to be important in cells that utilize this variant cell cycle.

## Materials and Methods

### Care of fly strains and crosses

Fly crosses were all carried out at 25° C with 50% humidity and a 12-hour light/dark cycle. Food (per liter) contained 18g yeast, 22g molasses, 80g malt extract, 9g agar, 65 cornmeal, 2.3g methyl para-benzoic acid, 6.35ml propionic acid. Sex of dissected larvae for imaginal disc studies were not determined. For Western blot and real-time PCR analyses, the number of male and female larvae were equal across the genotypes examined. For studies comparing wild-type and *kdm5* mutant larvae, we matched animals for developmental stage, and not chronological age, as we have done previously (Drelon et al., 2018). Thus, wild-type wandering 3^rd^ instar larvae were ∼120 hours after egg laying, while *kdm5*^*140*^ larvae were ∼10 days old.

### Real-time PCR

Total RNA was purified from either whole larvae or dissected brain-ring gland complexes using TRIzol. Reverse transcription was carried out using 1 μ g of RNA (or 5 μg of RNA for ecdysone biosynthetic genes) using a Verso cDNA kit (Thermo-Fisher AB1453A). Real-time PCR used the Power SYBR Green Master Mix and was performed in Applied Biosystems Step ONE plus real-time PCR system. Changes to gene expression were determined by normalizing samples to *rp49* (*RpL32*).

### Western blot

Western blots were carried out with dissected wing discs or ring glands. Samples were dissected in 1xPBS and transferred to 1xNuPAGE LDS sample buffer, run on a 4-12% Bis-Tris 1mm gel and transferred to PVDF. Secondary antibodies were donkey anti-mouse IgG 680RD or donkey anti-rabbit IgG 800CW. Blots were scanned using a LI-COR Odyssey Infrared scanner and quantified using LI-COR imaging software v3.0.

### Immunostaining of wing discs

Third instar wing imaginal discs were dissected in 1xPBS and fixed in 4% paraformaldehyde for 30 minutes. After blocking in 0.1% BSA/TBST (1xPBS, 0.2% Triton, 0.1% BSA) for 1 hour, samples were incubated with anti-Dcp-1 overnight at 4°C. Wing discs were washed with PBST and incubated with anti-rabbit Alexa Fluro 568 secondary antibody for 2 hours at 4°C. Samples were mounted in Vectashield for microscopy.

### Edu incorporation into prothoracic glands

For EdU staining, click-IT Edu kit and Alexa Fluor 594 Azide were used. Third instar larvae were dissected in Schneider’s *Drosophila* medium and brain-ring gland complexes were in incubated in 50μM EdU in Schneider’s Drosophila medium for 2.5 hours at room temperature. After washing in 1xPBS, tissues were fixed in 4% paraformaldehyde for 30 minutes. Tissues were blocked in 3% BSA for 30 minutes and then permeabilized in 0.5% triton and detected using 1x click-IT reaction buffer, CuSO4, Alexa Fluor 594 Azide and reaction buffer additive for 30 minutes. DNA was stained using 4′, 6-diamidino-2-phenylindole (DAPI) and brain-ring gland complexes were mounted in Vectashield for microscopy.

### Larval feeding of 20E

Control wild-type or *kdm5*^*140*^/CyO-GFP flies were allowed to mate and lay eggs for 6 hours. At 96 hours AEL, a mixture of dry yeast, 20-hydroxyecdysone (0.33mg/ml in 100% ethanol) or control ethanol alone in 500µl of ddH_2_O was added to each vial. *kdm5*^*140*^ homozygous larvae or pupae were identified by the absence of the CyO-GFP balancer.

### 20E quantification

Ecdysone was extracted from whole larvae as previously described (Moeller et al., 2017). Briefly, ten 3^rd^ instar larvae were washed in ddH_2_O and then homogenized in 0.5ml methanol. After centrifugation, the pellet was re-extracted in 0.5ml methanol and then in 0.5ml ethanol. The three supernatants were mixed and 0.5ml was evaporated using a SpeedVac. The pellet was dissolved in 100µl of EIA buffer and subjected to the 20-hydroxyecdysone EIA Kit.

### Lifespan analyses

Lifespan analyses was carried out by collecting adult flies 24-36 hours after eclosion and placing flies into vials at a density of 20 animals or fewer. Survival was quantified by transferring flies every two days into vials with fresh food and counting the number of dead animals. Survival analyses were carried out using Log-rank (Mantel-Cox) test in Prism v8.0.

### Developmental delay quantification

Female and male flies were placed in a vial and allowed to lay eggs for 16 hours. Starting at day 4 AEL, the number of animals that had pupariated were scored every 24 hours. Pupal genotype was ascertained by balancing *kdm5*^*140*^ using a CyO-GFP chromosome.

### Fertility analyses

To determine fertility, individual female flies were placed in a vial with 5 wild-type male flies and individual males were placed in a vial with 3-5 virgin wild-type females and allowed to mate and lay eggs for 4 days. If larvae were present in the vial by day 5, the fly was designated fertile. In cases where the fly died before day 3, we were unable confidently assess fertility and these animals were eliminated from the analyses.

### Image acquisition and processing

Adult fly images were obtained using Zeiss Discovery.V12 SteREO or ZEISS ApoTome microscopes and captured using AxioVision Release 4.8 software. Images of larval tissues were obtained using Zeiss AxioImager.M12 microscope and AxioVision SE64 Release 4.9.1 software. All images were processed using Adobe Photoshop CC 2019.

### Statistical analyses

All experiments were done in biological triplicate (minimum) and Ns are provided for each experiment. Fisher’s exact test was carried out in R. Student’s t-test, one-way ANOVA and Wilcoxon rank-sum tests were carried out using GraphPad Prism.

### Resource information

| REAGENT | SOURCE | IDENTIFIER |
| --- | --- | --- |
| Antibodies |  |  |
| KDM5 | Secombe lab | (Secombe et al., 2007) |
| Alpha-tubulin | DSHB | 12G10; 1:2000 |
| Phosphorylated ERK | Cell Signaling | 9106; 1:250 |
| ERK | Cell Signaling | 39763; 1:250 |
| Histone H3 | Active Motif | 39763; 1:1000 |
| S10 phosphorylated histone H3 | Cell Signaling | 9701S; 1:1000 |
| Atk | Cell Signaling | 9272; 1:200 |
| Dcp-1 | Cell Signaling | 9578; 1:100 |
| Ser505 phosphorylated Akt | Cell Signaling | 4054S; 1:200 |
| Donkey anti-mouse IgG 680RD | LI-COR | 925-68072; 1:8000 |
| Donkey anti-rabbit IgG 800CW | LI-COR | 926-32213; 1:8000 |
| Alexa Fluor 568 anti-rabbit | Invitrogen | A11036 |
| Alexa Fluor 594 Azide | Invitrogen | A10270 |
| Chemicals, Peptides, and Recombinant Proteins |  |  |
| Paraformaldehyde | Fisher Scientific | 28906 |
| Schneider's Drosophila medium | Fisher Scientific | 21720024 |
| 20-hydroxyecdysone | Fisher Scientific | H5142 |
| 10xPBS | Fisher Scientific | 70011069 |
| Trizol | Invitrogen | 15596026 |
| Vectashield | Vector Labs | H-1000 |
| Critical Commercial Assays |  |  |
| Click-IT EdU | Invitrogen | C10337 |
| 20-hydroxyecdysone EIA kit | Cayman Chemical | 501390 |
| Verso cDNA synthesis kit | Fisher-Thermo | AB-1453 |
| Power SYBR green master mix (real-time PCR) | Fisher-Thermo | 4367659 |
| 1xNuPAGE LDS sample buffer | Invitrogen | NP0008 |
| 4-12% Bis-Tris 1mm gels | Invitrogen | NP0322 |
| PVDF | Fisher Scientific | 88518 |
| Experimental Models: Organisms/Strains |  |  |
| *kdm5<sup>140</sup>* | Secombe lab | (Drelon et al., 2018) |
| Wild type | *kdm5<sup>140</sup> ; g[kdm5<sup>WT</sup>:HA]attP-86Fb* | (Zamurad et al., 2018) |
| *w<sup>1118</sup>* | Bloomington Drosophila Stock Center (BDSC) | BL5905 |
| UASp-*kdm5* | Secombe lab | (Secombe et al., 2007) |
| Ubiquitin-Gal4 | BDSC | BL32251 |
| 32B-Gal4 | BDSC | BL1782 |
| T80-Gal4 | BDSC | BL1878 |
| Mai60-Gal4 | BDSC | BL30811 |
| spok-Gal4 | Michael O'Connor | (Shimell et al., 2018) |
| phm-Gal4 | Michael O'Connor | (Rewitz et al., 2009) |
| Aug21-Gal4 | BDSC | BL20137 |
| Akh-Gal4 | BDSC | BL25684 |
| dilp2-Gal4 | BDSC | BL37516 |
| CG-Gal4 | BDSC | BL7011 |
| Mef2-Gal4 | BDSC | BL27390 |
| UAS-torso | Michael O'Connor | (Moeller et al., 2017) |
| UAS-cD8GFP | BDSC | BL8768 |
| UAS-*cyclin E* | Helena Richardson | (Richardson et al., 1995) |
| Software and Algorithms |  |  |
| GraphPad v8.0.2 | GraphPad Software | N/A |
| R v3.3.2 | http://www.R-project.org/ | N/A |

### Primers

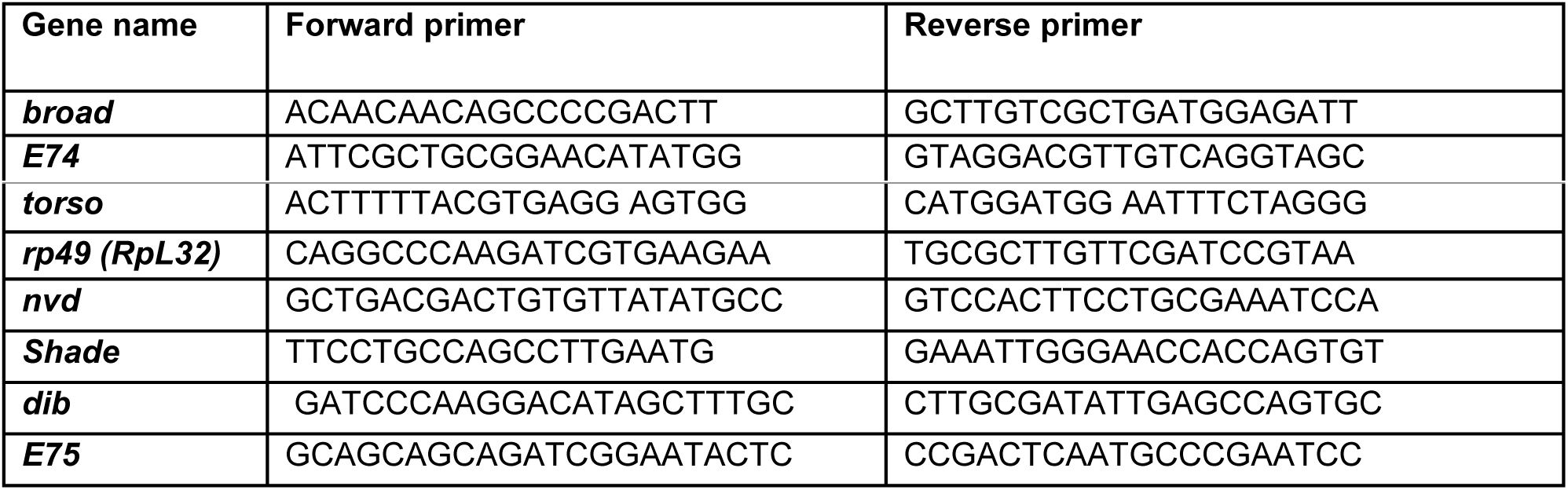

## Acknowledgements

We would like to thank members of the Secombe lab and Nick Baker’s lab for insights at all stages of this project. We are very grateful for fly strains donated by Michael O’Connor and Helena Richardson in addition to the Bloomington Drosophila Stock Center (NIH P400D018537). The 12G10 monoclonal antibody was obtained from the Developmental Studies Hybridoma bank, created by the NICHD of the NIH and maintained at The University of Iowa. We also thank the NIH (R01 GM112783) and the Einstein Cancer Center Support Grant P30 CA013330.

## Author Contribution

Conceptualization, J.S., C.D. and H.B.; Methodology C.D, and H.B.; Investigation, C.D. and H.B.; Writing – original draft, J.S., Writing – Reviewing and Editing, J.S., C.D. and H.B.; Funding acquisition, J.S., Supervision, J.S.

## Declaration of Interests

The authors declare no competing interests.

## Bibliography

Accari, S. L. & Fisher, P. R. 2015. Emerging Roles of JmjC Domain-Containing Proteins. Int Rev Cell Mol Biol, 319, 165–220.

Aggarwal, S. K. & King, R. C. 1969. A comparative study of the ring glands from wild type and 1(2)gl mutant Drosophila melanogaster. J Morphol, 129, 171–99.

Aguila, J. R., Suszko, J., Gibbs, A. G. & Hoshizaki, D. K. 2007. The role of larval fat cells in adult Drosophila melanogaster. J Exp Biol, 210, 956–63.

Albert, M., Schmitz, S. U., Kooistra, S. M., Malatesta, M., Morales torres, C., Rekling, J. C., Johansen, J. V., Abarrategui, I. & Helin, K. 2013. The histone demethylase Jarid1b ensures faithful mouse development by protecting developmental genes from aberrant H3K4me3. PLoS Genet, 9, e1003461.

Bannister, A. J. & Kouzarides, T. 2011. Regulation of chromatin by histone modifications. Cell Res, 21, 381–95.

Barrett, A., Santangelo, S., Tan, K., Catchpole, S., Roberts, K., Spencer-dene, B., Hall, D., Scibetta, A., Burchell, J., Verdin, E., Freemont, P. & Taylor-papadimitriou, J. 2007. Breast cancer associated transcriptional repressor PLU-1/JARID1B interacts directly with histone deacetylases. International Journal of Cancer, 121, 265–275.

Benayoun, B. A., Pollina, E. A., Ucar, D., Mahmoudi, S., Karra, K., Wong, E. D., Devarajan, K., Daugherty, A. C., Kundaje, A. B., Mancini, E., Hitz, B. C., Gupta, R., Rando, T. A., Baker, J. C., Snyder, M. P., Cherry, J. M. & Brunet, A. 2014. H3K4me3 breadth is linked to cell identity and transcriptional consistency. Cell, 158, 673–88.

Blair, L. P., Cao, J., Zou, M. R., Sayegh, J. & Yan, Q. 2011. Epigenetic Regulation by Lysine Demethylase 5 (KDM5) Enzymes in Cancer. Cancers (Basel), 3, 1383–1404.

Brookes, E., Laurent, B., Ounap, K., Carroll, R., Moeschler, J. B., Field, M., Schwartz, C. E., Gecz, J. & Shi, Y. 2015. Mutations in the intellectual disability gene KDM5C reduce protein stability and demethylase activity. Hum Mol Genet, 24, 2861–72.

Cao, J., Liu, Z., Cheung, W. K., Zhao, M., Chen, S. Y., Chan, S. W., Booth, C. J., Nguyen, D. X. & Yan, Q. 2014. Histone demethylase RBP2 is critical for breast cancer progression and metastasis. Cell Rep, 6, 868–77.

Catchpole, S., Spencer-dene, B., Hall, D., Santangelo, S., Rosewell, I., Guenatri, M., Beatson, R., Scibetta, A. G., Burchell, J. M. & Taylor-papadimitriou, J. 2011. PLU-1/JARID1B/KDM5B is required for embryonic survival and contributes to cell proliferation in the mammary gland and in ER+ breast cancer cells. Int J Oncol, 38, 1267–77.

Chen, K., Luan, X., Liu, Q., Wang, J., Chang, X., Snijders, A. M., Mao, J. H., Secombe, J., Dan, Z., Chen, J. H., Wang, Z., Dong, X., Qiu, C., Chang, X., Zhang, D., Celniker, S. E. & Liu, X. 2019. Drosophila Histone Demethylase KDM5 Regulates Social Behavior through Immune Control and Gut Microbiota Maintenance. Cell Host Microbe, 25, 537–552 e8.

Christesen, D., Yang, Y. T., Somers, J., Robin, C., Sztal, T., Batterham, P. & Perry, T. 2017. Transcriptome Analysis of Drosophila melanogaster Third Instar Larval Ring Glands Points to Novel Functions and Uncovers a Cytochrome p450 Required for Development. G3 (Bethesda), 7, 467–479.

Collins, B. E., Greer, C. B., Coleman, B. C. & Sweatt, J. D. 2019. Histone H3 lysine K4 methylation and its role in learning and memory. Epigenetics Chromatin, 12, 7.

Cong, B., Ohsawa, S. & Igaki, T. 2018. JNK and Yorkie drive tumor progression by generating polyploid giant cells in Drosophila. Oncogene, 37, 3088–3097.

Danielsen, E. T., Moeller, M. E. & Rewitz, K. F. 2013. Nutrient signaling and developmental timing of maturation. Curr Top Dev Biol, 105, 37–67.

Danielsen, E. T., Moeller, M. E., Yamanaka, N., Ou, Q., Laursen, J. M., Soenderholm, C., Zhuo, R., Phelps, B., Tang, K., Zeng, J., Kondo, S., Nielsen, C. H., Harvald, E. B., Faergeman, N. J., Haley, M. J., O’connor, K. A., King-jones, K., O’connor, M. B. & Rewitz, K. F. 2016. A Drosophila Genome-Wide Screen Identifies Regulators of Steroid Hormone Production and Developmental Timing. Dev Cell, 37, 558–70.

Davoli, T. & De lange, T. 2011. The causes and consequences of polyploidy in normal development and cancer. Annu Rev Cell Dev Biol, 27, 585–610.

Drelon, C., Belalcazar, H. M. & Secombe, J. 2018. The Histone Demethylase KDM5 Is Essential for Larval Growth in Drosophila. Genetics.

Fox, D. T. & Duronio, R. J. 2013. Endoreplication and polyploidy: insights into development and disease. Development, 140, 3–12.

Frawley, L. E. & Orr-weaver, T. L. 2015. Polyploidy. Curr Biol, 25, R353–8.

Gajan, A., Barnes, V. L., Liu, M., Saha, N. & Pile, L. A. 2016. The histone demethylase dKDM5/LID interacts with the SIN3 histone deacetylase complex and shares functional similarities with SIN3. Epigenetics Chromatin, 9, 4.

Gaziova, I., Bonnette, P. C., Henrich, V. C. & Jindra, M. 2004. Cell-autonomous roles of the ecdysoneless gene in Drosophila development and oogenesis. Development, 131, 2715–25.

Gilbert, L. I. 2004. Halloween genes encode P450 enzymes that mediate steroid hormone biosynthesis in Drosophila melanogaster. Mol Cell Endocrinol, 215, 1–10.

Gildea, J. J., Lopez, R. & Shearn, A. 2000. A screen for new trithorax group genes identified little imaginal discs, the Drosophila melanogaster homologue of human retinoblastoma binding protein 2. Genetics, 156, 645–663.

Gu, S. H., Young, S. C., Lin, J. L. & Lin, P. L. 2011. Involvement of PI3K/Akt signaling in PTTH-stimulated ecdysteroidogenesis by prothoracic glands of the silkworm, Bombyx mori. Insect Biochem Mol Biol, 41, 197–202.

Han, M., Xu, W., Cheng, P., Jin, H. & Wang, X. 2017. Histone demethylase lysine demethylase 5B in development and cancer. Oncotarget, 8, 8980–8991.

Howe, F. S., Fischl, H., Murray, S. C. & Mellor, J. 2017. Is H3K4me3 instructive for transcription activation? Bioessays, 39, 1–12.

Hrdlicka, L., Gibson, M., Kiger, A., Micchelli, C., Schober, M., Schock, F. & Perrimon, N. 2002. Analysis of twenty-four Gal4 lines in Drosophila melanogaster. Genesis, 34, 51–7.

Iwase, S., Brookes, E., Agarwal, S., Badeaux, A. I., Ito, H., Vallianatos, C. N., Tomassy, G. S., Kasza, T., Lin, G., Thompson, A., Gu, L., Kwan, K. Y., Chen, C., Sartor, M. A., Egan, B., Xu, J. & Shi, Y. 2016. A Mouse Model of X-linked Intellectual Disability Associated with Impaired Removal of Histone Methylation. Cell Rep, 14, 1000–9.

Iwase, S., Lan, F., Bayliss, P., De la torre-ubieta, L., Huarte, M., Qi, H. H., Whetstine, J. R., Bonni, A., Roberts, T. M. & Shi, Y. 2007. The X-linked mental retardation gene SMCX/JARID1C defines a family of histone H3 lysine 4 demethylases. Cell, 128, 1077–1088.

Kim, J. H., Lee, J. H., Lee, I. S., Lee, S. B. & Cho, K. S. 2017. Histone Lysine Methylation and Neurodevelopmental Disorders. Int J Mol Sci, 18.

Klose, R. J., Yan, Q., Tothova, Z., Yamane, K., Erdjument-bromage, H., Tempst, P., Gilliland, D. G., Zhang, Y. & Kaelin, W. G. J. 2007. The Retinoblastoma binding protein RBP2 is a H3K4 demethylase. Cell, 128, 889–900.

Lee, G. & Park, J. H. 2004. Hemolymph sugar homeostasis and starvation-induced hyperactivity affected by genetic manipulations of the adipokinetic hormone-encoding gene in Drosophila melanogaster. Genetics, 167, 311–23.

Lee, N., Erdjument-bromage, H., Tempst, P., Jones, R. S. & Zhang, Y. 2009. The H3K4 Demethylase Lid Associates with and Inhibits the Histone Deacetylase Rpd3. Molecular and Cellular Biology, 29, 1401–1410.

Li, L., Greer, C., Eisenman, R. N. & Secombe, J. 2010. Essential functions of the histone demethylase lid. PLoS Genet, 6, e1001221.

Liu, X., Greer, C. & Secombe, J. 2014. KDM5 interacts with Foxo to modulate cellular levels of oxidative stress. PLoS Genet, 10, e1004676.

Liu, X. & Secombe, J. 2015. The Histone Demethylase KDM5 Activates Gene Expression by Recognizing Chromatin Context through Its PHD Reader Motif. Cell Rep, 13, 2219–31.

Lloret-llinares, M., Perez-lluch, S., Rossell, D., Moran, T., Ponsa-cobas, J., Auer, H., Corominas, M. & Azorin, F. 2012. dKDM5/LID regulates H3K4me3 dynamics at the transcription-start site (TSS) of actively transcribed developmental genes. Nucleic Acids Research.

Mariani, L., Lussi, Y. C., Vandamme, J., Riveiro, A. & Salcini, A. E. 2016. The H3K4me3/2 histone demethylase RBR-2 controls axon guidance by repressing the actin-remodeling gene wsp-1. Development, 143, 851–63.

Martin, H. C., Jones, W. D., Mcintyre, R., Sanchez-andrade, G., Sanderson, M., Stephenson, J. D., Jones, C. P., Handsaker, J., Gallone, G., Bruntraeger, M., Mcrae, J. F., Prigmore, E., Short, P., Niemi, M., Kaplanis, J., Radford, E. J., Akawi, N., Balasubramanian, M., Dean, J., Horton, R., Hulbert, A., Johnson, D. S., Johnson, K., Kumar, D., Lynch, S. A., Mehta, S. G., Morton, J., Parker, M. J., Splitt, M., Turnpenny, P. D., Vasudevan, P. C., Wright, M., Bassett, A., Gerety, S. S., Wright, C. F., Fitzpatrick, D. R., Firth, H. V., Hurles, M. E., Barrett, J. C. & DECIPHERING DEVELOPMENTAL DISORDERS, S. 2018. Quantifying the contribution of recessive coding variation to developmental disorders. Science.

Mcbrayer, Z., Ono, H., Shimell, M., Parvy, J. P., Beckstead, R. B., Warren, J. T., Thummel, C. S., Dauphin-villemant, C., Gilbert, L. I. & O’connor, M. B. 2007. Prothoracicotropic hormone regulates developmental timing and body size in Drosophila. Dev Cell, 13, 857–71.

Mccann, T. S., Sobral, L. M., Self, C., Hsieh, J., Sechler, M. & Jedlicka, P. 2019. Biology and targeting of the Jumonji-domain histone demethylase family in childhood neoplasia: a preclinical overview. Expert Opin Ther Targets, 1–14.

Mirabella, A. C., Foster, B. M. & Bartke, T. 2016. Chromatin deregulation in disease. Chromosoma, 125, 75–93.

Moeller, M. E., Nagy, S., Gerlach, S. U., Soegaard, K. C., Danielsen, E. T., Texada, M. J. & Rewitz, K. F. 2017. Warts Signaling Controls Organ and Body Growth through Regulation of Ecdysone. Curr Biol, 27, 1652–1659 e4.

Moran, T., Bernues, J. & Azorin, F. 2015. The Drosophila histone demethylase dKDM5/LID regulates hematopoietic development. Dev Biol, 405, 260–8.

Moshkin, Y. M., Kan, T. W., Goodfellow, H., Bezstarosti, K., Maeda, R. K., Pilyugin, M., Karch, F., Bray, S. J., Demmers, J. A. & Verrijzer, C. P. 2009. Histone chaperones ASF1 and NAP1 differentially modulate removal of active histone marks by LID-RPD3 complexes during NOTCH silencing. Molecular Cell, 35, 782–93.

Navarro-costa, P., Mccarthy, A., Prudencio, P., Greer, C., Guilgur, L. G., Becker, J. D., Secombe, J., Rangan, P. & Martinho, R. G. 2016. Early programming of the oocyte epigenome temporally controls late prophase I transcription and chromatin remodelling. Nat Commun, 7, 12331.

Nishibuchi, G., Shibata, Y., Hayakawa, T., Hayakawa, N., Ohtani, Y., Sinmyozu, K., Tagami, H. & Nakayama, J. 2014. Physical and functional interactions between the histone H3K4 demethylase KDM5A and the nucleosome remodeling and deacetylase (NuRD) complex. J Biol Chem, 289, 28956–70.

Niwa, R. & Niwa, Y. S. 2014. Enzymes for ecdysteroid biosynthesis: their biological functions in insects and beyond. Biosci Biotechnol Biochem, 78, 1283–92.

Ohhara, Y., Kobayashi, S. & Yamanaka, N. 2017. Nutrient-Dependent Endocycling in Steroidogenic Tissue Dictates Timing of Metamorphosis in Drosophila melanogaster. PLoS Genet, 13, e1006583.

Orr-weaver, T. L. 2015. When bigger is better: the role of polyploidy in organogenesis. Trends Genet, 31, 307–15.

Paroni, G., Bolis, M., Zanetti, A., Ubezio, P., Helin, K., Staller, P., Gerlach, L. O., Fratelli, M., Neve, R. M., Terao, M. & Garattini, E. 2018. HER2-positive breast-cancer cell lines are sensitive to KDM5 inhibition: definition of a gene-expression model for the selection of sensitive cases. Oncogene.

Plch, J., Hrabeta, J. & Eckschlager, T. 2019. KDM5 demethylases and their role in cancer cell chemoresistance. Int J Cancer, 144, 221–231.

Rewitz, K. F., Yamanaka, N., Gilbert, L. I. & O’connor, M. B. 2009. The insect neuropeptide PTTH activates receptor tyrosine kinase torso to initiate metamorphosis. Science, 326, 1403–5.

Richardson, H., O’keefe, L. V., Marty, T. & Saint, R. 1995. Ectopic cyclin E expression induces premature entry into S phase and disrupts pattern formation in the Drosophila eye imaginal disc. Development, 121, 3371–9.

Roesch, A., Vultur, A., Bogeski, I., Wang, H., Zimmermann, K. M., Speicher, D., Korbel, C., Laschke, M. W., Gimotty, P. A., Philipp, S. E., Krause, E., Patzold, S., Villanueva, J., Krepler, C., Fukunaga-kalabis, M., Hoth, M., Bastian, B. C., Vogt, T. & Herlyn, M. 2013. Overcoming Intrinsic Multidrug Resistance in Melanoma by Blocking the Mitochondrial Respiratory Chain of Slow-Cycling JARID1B(high) Cells. Cancer Cell, 23, 811–25.

Santos-rosa, H., Schneider, R., Bannister, A. J., Sherriff, J., Bernstein, B. E., Emre, N. C. T., Schreiber, S. L., Mellor, J. & Kouzarides, T. 2002. Active genes are tri-methylated at K4 of histone H3. Nature, 419, 407–411.

Saxton, R. A. & Sabatini, D. M. 2017a. mTOR Signaling in Growth, Metabolism, and Disease. Cell, 168, 960–976.

Saxton, R. A. & Sabatini, D. M. 2017b. mTOR Signaling in Growth, Metabolism, and Disease. Cell, 169, 361–371.

Secombe, J., Li, L., Carlos, L. S. & Eisenman, R. N. 2007. The Trithorax group protein Lid is a trimethyl histone H3K4 demethylase required for dMyc-induced cell growth. Genes & Development, 21, 537–551.

Sharma, K., Singh, J., Frost, E. E. & Pillai, P. P. 2018. MeCP2 in central nervous system glial cells: current updates. Acta Neurobiol Exp (Wars), 78, 30–40.

Shimell, M., Pan, X., Martin, F. A., Ghosh, A. C., Leopold, P., O’connor, M. B. & Romero, N. M. 2018. Prothoracicotropic hormone modulates environmental adaptive plasticity through the control of developmental timing. Development, 145.

Tahiliani, M., Mei, P. C., Fang, R., Leonor, T., Rutenberg, M., Shimizu, F., Li, J., Rao, A. & Shi, Y. J. 2007. The histone H3K4 demethylase SMCX links REST target genes to X-linked mental retardation. Nature, 447, 601-+.

Tarayrah, L., Li, Y., Gan, Q. & Chen, X. 2015. Epigenetic regulator Lid maintains germline stem cells through regulating JAK-STAT signaling pathway activity. Biol Open, 4, 1518–27.

Unhavaithaya, Y. & Orr-weaver, T. L. 2012. Polyploidization of glia in neural development links tissue growth to blood-brain barrier integrity. Genes Dev, 26, 31–6.

Vallianatos, C. N., Farrehi, C., Friez, M. J., Burmeister, M., Keegan, C. E. & Iwase, S. 2018. Altered Gene-Regulatory Function of KDM5C by a Novel Mutation Associated With Autism and Intellectual Disability. Front Mol Neurosci, 11, 104.

Vallianatos, C. N. & Iwase, S. 2015. Disrupted intricacy of histone H3K4 methylation in neurodevelopmental disorders. Epigenomics, 7, 503–19.

Xhabija, B. & Kidder, B. L. 2018. KDM5B is a master regulator of the H3K4-methylome in stem cells, development and cancer. Semin Cancer Biol.

Xiang, J., Bandura, J., Zhang, P., Jin, Y., Reuter, H. & Edgar, B. A. 2017. EGFR-dependent TOR-independent endocycles support Drosophila gut epithelial regeneration. Nat Commun, 8, 15125.

Yamamoto, S., Wu, Z., Russnes, H. G., Takagi, S., Peluffo, G., Vaske, C., Zhao, X., Moen vollan, H. K., Maruyama, R., Ekram, M. B., Sun, H., Kim, J. H., Carver, K., Zucca, M., Feng, J., Almendro, V., Bessarabova, M., Rueda, O. M., Nikolsky, Y., Caldas, C., Liu, X. S. & Polyak, K. 2014. JARID1B is a luminal lineage-driving oncogene in breast cancer. Cancer Cell, 25, 762–77.

Yamanaka, N., Rewitz, K. F. & O’connor, M. B. 2013. Ecdysone control of developmental transitions: lessons from Drosophila research. Annu Rev Entomol, 58, 497–516.

Yamane, K., Tateishi, K., Klose, R. J., Fang, J., Fabrizio, L. A., Erdjument-bromage, H., Taylor-papadimitriou, J., Tempst, P. & Zhang, Y. 2007. PLU-1 is a H3K4 demethylase involved in transcriptional repression and breast cancer cell proliferation. Molecular Cell, 25, 801–812.

Zamurrad, S., Hatch, H. A. M., Drelon, C., Belalcazar, H. M. & Secombe, J. 2018. A Drosophila Model of Intellectual Disability Caused by Mutations in the Histone Demethylase KDM5. Cell Rep, 22, 2359–2369.

Zhaunova, L., Ohkura, H. & Breuer, M. 2016. Kdm5/Lid Regulates Chromosome Architecture in Meiotic Prophase I Independently of Its Histone Demethylase Activity. PLoS Genet, 12, e1006241.

